# Integrated phenotypic and proteomic screening identifies top-tier Alzheimer’s disease therapeutic targets

**DOI:** 10.1101/2025.10.16.682926

**Authors:** Gregory A Cary, Qianjin Li, Jesse C Wiley, Carolyn A Paisie, Yuhong Du, Elizabeth L Zoeller, Duc Duong, Haian Fu, Nicholas T. Seyfried, Allan I. Levey, Ranjita Betarbet, Gregory W Carter, The Emory-Sage-SGC-JAX TREAT-AD Center

**Author notes:** **Corresponding author:** Gregory W Carter, The Jackson Laboratory, Bar Harbor ME. These authors contributed equally (Gregory A Cary, Qianjin Li).

## Abstract

**Introduction:** Alzheimer’s disease (AD) is a complex neurodegenerative disorder. Hundreds of therapeutic targets have been nominated through genetic and multi-omic studies, but effective prioritization remains a major bottleneck.

**Methods:** We applied an integrative screening framework to assess 29 candidate targets from risk-enriched biological domains. Using disease-relevant murine BV2 microglial cell lines with stable Psen2 knockdown, we performed siRNA-mediated perturbations followed by cellular phenotypic assays and quantitative proteomics.

**Results:** Twenty-five candidate targets significantly altered at least one phenotype, with stronger effects in Psen2 knockdown cells. Integrated proteomic analyses identified several perturbations that reversed AD-associated molecular patterns. Five targets—Ap2a2, Pdhb, Pdha1, Dlat, and Psmc3— impacted both phenotypes and related proteomic responses.

**Discussion:** We established a scalable platform for target functional validation that bridges unbiased systems-level assessments of AD risk with experimental evidence. The ESSJ TREAT-AD center will prioritize further resource development for these validated targets.

## 1. BACKGROUND

Alzheimer’s disease (AD) is a late-onset, progressive neurodegenerative disorder that presents an immense and escalating burden on global healthcare systems, patients, and caregivers [1]. There is an urgent need for the development of novel therapeutic interventions that can ameliorate these burdens. However, therapeutic development efforts are challenging given the complex, multifactorial nature of the disease. The interplay among genetic, lifestyle, and environmental risk factors over decades of life obscures which disease-altered processes are optimal targets for effective interventions [2]. Systems-level analyses such as genome-wide association studies (GWAS) and multi-omic analyses have been used to address and catalog this complexity. These studies have yielded extensive lists of potential therapeutic targets. The largest genome wide association studies implicate over 75 genomic loci in disease risk [3], while the AD Knowledge Portal (agora.adknowledgeportal.org) currently lists over 900 targets nominated for therapeutic development based on the Accelerating Medicines Partnership for Alzheimer’s Disease (AMP-AD) nominations [4]. Effectively prioritizing these candidate targets remains a significant bottleneck.

We have previously described an approach to integrate signatures of risk from AD case data [5]. This framework uses evidence from genetic association studies (e.g. GWAS, QTL) and from transcriptomic and proteomic assessment of differential expression in postmortem brain to score AD risk genome-wide. This risk score is paired with the AD biological domains (biodomains), which enumerate the molecular features commonly implicated in the disease. The biodomains are functionalized via annotation of each with a set of largely unique Gene Ontology (GO) terms. Using these resources highlighted especially strong enrichment for disease risk among the Synapse, Immune Response, Lipid Metabolism, and Mitochondrial Metabolism domains [5]. We interrogated these risk-enriched processes with orthogonal approaches to articulate interpretable hypotheses based on the systems-level signatures of risk. We generated two exemplar hypotheses with this approach: (1) mitochondrial hypometabolism leads to an increased susceptibility to neurodegeneration, and (2) chronic neuroinflammation via over-activation of the innate immune response leads to neurodegenerative pathology in AD. These hypotheses are not novel and reflect consensus opinions from recent years regarding disease pathogenesis [6–8]. However, these hypotheses have been developed through systematic integration of disease risk signatures, and bioinformatic pipelines are used to identify candidate risk-enriched targets related to each overarching hypothesis area. Each hypothesis emanates from a specific GO term (i.e. “mitochondrion” GO:0005739 and “innate immune response” GO:0045087), each having several hundred annotated risk genes. It is not clear from the available data which of the predicted driver genes are capable of affecting the targeted biology in appropriate cellular contexts.

To resolve this question, we have developed a robust experimental system to screen candidate driver genes for those that are capable of affecting the biology under investigation (Figure 1). Given the nature of the exemplar hypotheses, we first used a microglial surrogate cell system to assess the impact of targeting the nominated genes. The genes were knocked down in the microglial surrogate lines and phenotypes relevant to mitochondrial function and immune response were assayed. Proteomic responses were also assessed following target knockdown. We integrated the results from cellular phenotype assays with proteomic responses to identify targets that both affect the phenotypes under investigation and impact associated biodomain terms in the proteomic results. Of the 29 candidate targets screened, we identify strong support for five targets - Ap2a2, Pdhb, Pdha1, Dlat, and Psmc3 - that both significantly impact the cellular phenotypes screened and corresponding biodomain terms. Importantly, perturbation of each of these targets demonstrated the capacity to reverse the proteomic signatures associated with disease in these experiments. This work establishes a platform to filter and validate prioritized candidate therapeutic targets to those that have the greatest potential to impact disease-relevant biology. This platform will be used by the Emory-Sage-SGC-JAX TREAT-AD center to prioritize candidate targets for the development of additional target enabling packages by the center.

**Figure 1.**
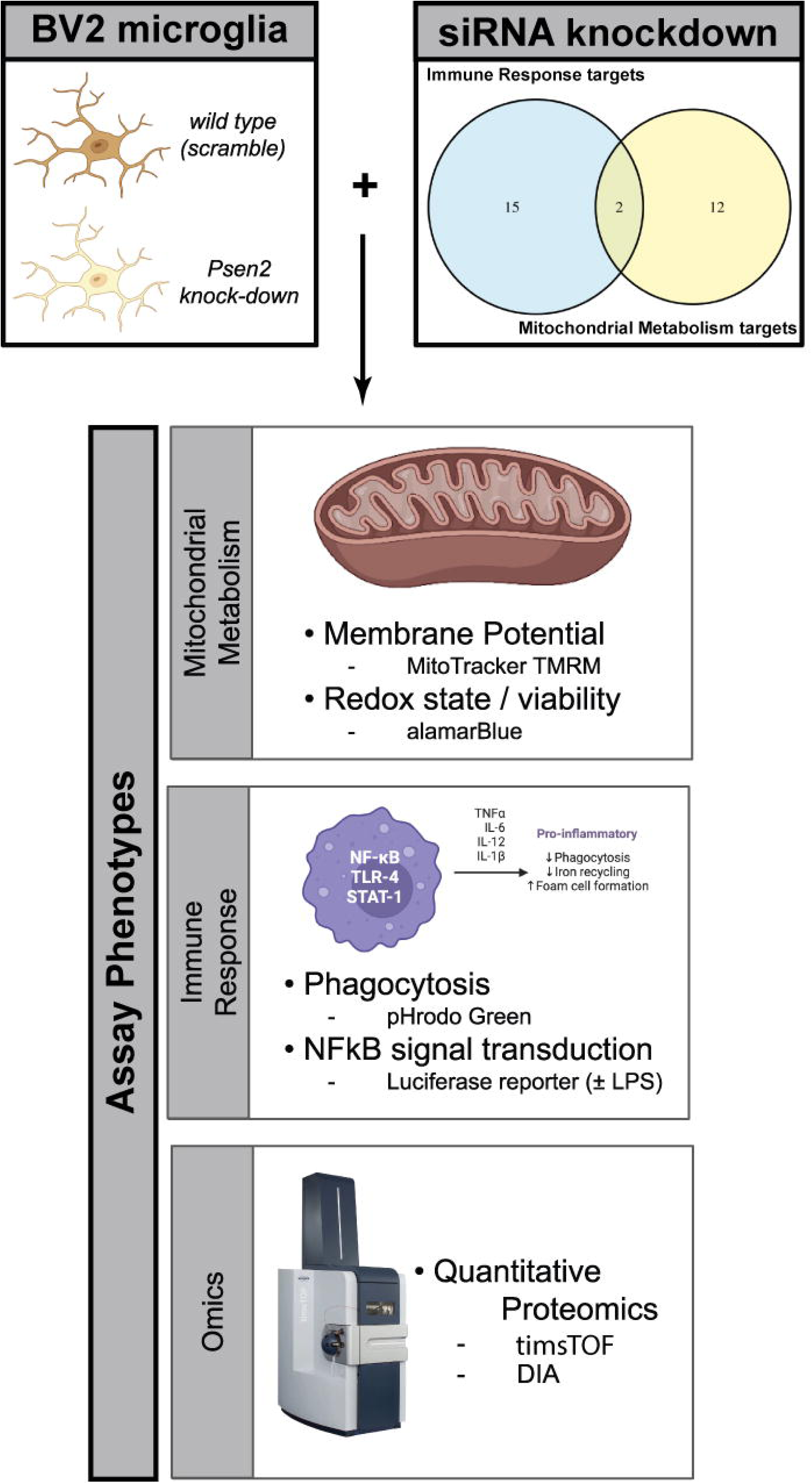
Overview of the design of the current study. siRNA knockdown was performed against 29 targets from two hypothesis areas (top right) in two different murine BV2 microglial cell lines — one wild-type (i.e., scramble) and one where Psen2 has been stably knocked down (top left). Following 72 hours of target knockdown, a series of cellular and molecular phenotypes were assayed including: MitoTracker TMRM and alamarBlue to assess mitochondrial function, and pHrodo Green and NFkB luciferase reporter activity. Quantitative proteomic measurements were also made following target knockdown. Created in BioRender. Cary, G. (2025) https://BioRender.com/scfycfr.

## 2. METHODS

### 2.1 Cell Culture and siRNA-Mediated Target Knockdown

Murine microglial BV2 cell lines stably expressing Presenilin 2 (Psen2) shiRNA or scrambled siRNA, as previously described [9], were kindly provided by Dr. Suman Jayadev (University of Washington School of Medicine in Seattle, Washington). Cells were cultured in DMEM/F12 (1:1) 10% FBS and 5 mg/ml puromycin selection antibiotic.

Candidate targets were selected from two broad hypothesis areas, each derived from related GO terms from the AD biological domains (biodomains) [5]: (1) immune response from the “innate immune response” term (GO:0045087) and (2) mitochondrial metabolism from the “mitochondrion” term (GO:0005739). These terms were both risk-enriched based on the results of GSEA using the AD target risk score [5] and were also implicated in early disease pseudotemporal states [10]. Through examination of protein-protein interaction networks centered on the proteins in each hypothesis area, integration with AMP-AD nominated targets from Agora (agora.adknowledgeportal.org), and a target novelty analysis, we identified 29 candidate targets — 17 targets within the Immune Response hypothesis area and 14 targets within the Mitochondrial Metabolism hypothesis area, with 2 common targets (Figure 1). Six of the 29 candidates have been previously nominated by AMP-AD investigators, and there is a mix of genes with transcripts and proteins that are both up and down in AD brains postmortem (Figure S1A). We confirmed the expression of the candidate target genes in publicly available gene expression datasets from BV2 cells (GEO accessions: GSE162526, GSE132739, and GSE137741) by averaging the log2 transformed expression values from each dataset and plotting a heatmap (Figure S1B).

A custom library of siRNAs against BioHyV targets (ON-TARGETplus™ siRNA) was purchased from Horizon Discovery, which consisted of a pool of four siRNA per target (Supplementary Table S1). Approximately 18–24 hours before transfection, cells were seeded in 384-well plates. 1200 cells in 25 μl media were dispensed into each well of a 384-well plate using multidrop combi (Thermo-Fisher Scientific). siRNAs were transfected with TransIT-X2 (Mirus, #MIR 6000) at 25 nM final concentration. After 72 hours of transfection, a panel of assays were performed as described in the following, including cell viability, mitochondrial membrane potential, phagocytosis, and NFκB reporter assays.

Knockdown efficiency of the siRNAs was confirmed with real-time PCR (Figure S1C). Cells were washed with PBS twice. Total RNA was isolated using the RNeasy Kit (Qiagen) following the protocol of the manufacturer. Complementary DNA was reverse-transcribed from 1 μg of total RNA in a 20-μl reaction with a reverse transcription kit (QIAGEN, #205313). The reverse transcription products were diluted 30 times with distilled H_2_O, and 1.5 μl was used as template for each real time PCR reaction. The reactions were performed using SsoAdvanced Universal SYBR® Green Supermix (BioRad, #1725274). The thermal cycling conditions were composed of an initial denaturation step at 95 °C for 1 min, 40 cycles at 95 °C for 10 s, and 60 °C for 50 s. The experiments were carried out in triplicate. Relative quantification of fold changes in gene expression was obtained by the 2^-ΔΔCt^ method, with GAPDH as the internal reference gene. The sequences of primers used for RT-PCR are listed in (Supplementary Table S2).

### 2.2 Cellular Phenotypic Assays

#### Cell viability assay

5 μl CellTiter-Blue reagent (Promega, #G8088) was added to each well of a 384-well plate after 72 hours after siRNA transfection. After incubating for 4 hours at 37°C, Fluorescence Intensity (FI) which is corresponding to the number of living cells was measured with PHERAstar FSX (BMG Labtech) plate reader (Ex. 560nm and Em. 590nm). Wells containing medium only were used as background signals. The assay was run in three separate batches with three biological replicates per batch for a total 9 measurements per siRNA and BV2 cell line.

#### Mitochondrial membrane potential assay

MitoTracker TMRM (ThermoFisher, #I34361) was diluted in culture medium containing the nuclear stain Hoechst 33342. 2 μl staining solution was added to each well to final concentration of 100 nM TMRM and 10 μg/ml Hoechst 33342. After 30 minutes incubation at 37°C, cells were imaged with ImageXpress® Micro (Molecular Devices) and the average fluorescence intensity of the cells were quantified based on cell number indicated by Hoechst staining with MetaXpress 6. The assay was run in seven separate batches with three biological replicates per batch for a total of 21 measurements per siRNA and BV2 cell line.

#### Phagocytosis assay

pHrodo Green Zymosan BioParticles Conjugate (ThermoFisher, # P35365) was dissolved in PBS and mixed with Hoechst 33342. 4 μg Zymosan Bioparticles Conjugates were added to each well with Hoechst 33342 at final concentration of 10 μg/ml. After 2 hr incubation at 37°C, cells were imaged with ImageXpress® Micro (Molecular Devices) and data were analyzed with MetaXpress 6. The assay was run in three separate batches with three to six biological replicates per batch for a total of 12 measurements per siRNA and BV2 cell line.

#### NF-kappaB reporter assay

Stable cell lines expressing luciferase driven by an NFκB response element were generated by transfecting BV2/Scramble and BV2/Psen2-KD cells with plasmid pGL4.32[luc2P/NFκB-RE/Hygro] (Promega, #E8491). 3 days after transfection, hygromycin was applied to cells at concentration of 400 μg/ml to select stable cell lines. For the NFκB reporter assay, cells harboring the luciferase reporter gene were transfected with siRNA for target gene knockdown.

3 days after transfection, cells were treated with either LPS (0.1 μg/ml) or vehicle for 6 hours. Luciferase signal was measured with EnVision 2103 Multilabel plate Reader (PekinElmer). The assay was run in three separate batches with four to five biological replicates per batch for a total of 14 measurements per siRNA, BV2 cell line, and LPS dose.

#### Statistical analyses

Each assay was analyzed by first computing an average across all biological replicates per batch, then a log2-fold change was computed by comparing the average target siRNA values to the average control siRNA values within the same BV2 cell line and LPS dose, where applicable. A linear mixed model with random effects was fit using the lmer function from the lme4 R package [11] to assess the statistical significance for each assay. The log2 fold change values were modeled as a combination of fixed effects (siRNA and BV2 cell line) and random effects (experimental batch). Calculation of estimated marginal means and post-hoc statistical testing was performed with the emmeans R package [12]. Assay hits were defined as any knockdown that achieved an effect with a Benjamini-Hochberg [13] corrected p-value ≤ 0.05. All results are reported (Supplemental Table S3).

### 2.3 Proteomic Analyses

#### Cell Homogenization and Protein Digestion

Cell pellets were lysed in lysis buffer (8 M urea, 10 mM Tris, 100 mM NaH2PO4, pH 8.5) and protein concentration was determined by bicinchoninic acid (BCA) assay (Pierce). For protein digestion, 25 μg of each sample was aliquoted and volumes normalized with additional lysis buffer. Samples were reduced with 5 mM dithiothreitol (DTT) at room temperature for 30 min, followed by 10 mM iodoacetamide (IAA) alkylation in the dark for another 30 min. Lysyl endopeptidase (Wako) at 1:25 (w/w) was added, and digestion allowed to proceed overnight. Samples were then 7-fold diluted with 50 mM ammonium bicarbonate. Trypsin (Promega) was then added at 1:25 (w/w) and digestion proceeded overnight. The peptide solutions were acidified to a final concentration of 1% (vol/vol) formic acid (FA) and 0.1% (vol/vol) trifluoroacetic acid (TFA) and desalted with a 10 mg HLB column (Oasis). Each HLB column was first rinsed with 1 mL of methanol, washed with 1 mL 50% (vol/vol) acetonitrile (ACN), and equilibrated with 2×1 mL 0.1% (vol/vol) TFA. The samples were then loaded onto the column and washed with 2×1 mL 0.1% (vol/vol) TFA. Elution was performed with 2 volumes of 0.5 mL 50% (vol/vol) ACN. From each sample, 10% of the eluent was split out for single shot DIA, 10% was used to make a combined global internal standard and 80% was saved for TMT labeling. All aliquots were dried by speedvac and stored.

#### Single Shot DIA Liquid Chromatography Mass Spectrometry

All samples were resuspended in 10 μl of loading buffer (0.1% FA, 0.03% TFA, 1% ACN) and 1 μl analyzed by liquid chromatography coupled to tandem mass spectrometry. Peptide eluents were separated on a custom in-house packed CSH 1.7 μm (15 cm × 150 μM internal diameter (ID) by a Ultimate U3000 RSLCnano (ThermoFisher Scientific). Buffer A was water with 0.1% (vol/vol) formic acid, and buffer B was 80% (vol/vol) acetonitrile in water with 0.1% (vol/vol) formic acid. Elution was performed over a 30 min gradient with flow rate at 1500 nL/min. The gradient was from 1% to 99% solvent B. Peptides were monitored on a timsTOF HT (Bruker Scientific) using the machine standard dia-PASEF method for short gradients (mass range from 475 to 1000, mobility range of 0.85 to 1.27 1/K0, 25 Da non-overlapping windows and a cycle time of 0.95 s).

#### Database Searches and Protein Quantification

All data were analyzed using DIA-NN (v1.8.1; [14]) searched against the UniProtKB mouse protein database (August 2020 with 91413 target sequences). The parameters were specified as follows: [CP1] canonical tryptic specificity, minimum fragment m/z of 200 and maximum fragment m/z of 5000, fixed modification for carbamidomethylation of cysteine (+57.02146 Da), variable modifications for methionine excision, oxidation of methionine (+15.994915 Da), and acetylation (+42.010565 Da), maximum of 2 missed cleavages, minimum peptide length of 7 and maximum peptide length of 50, minimum precursor m/z of 200 and maximum precursor m/z of 5000, minimum precursor charge of 1 and maximum precursor charge of 4, MS1 and MS2 mass accuracy of 10 ppm, smart profiling and MBR (match-between-runs) enabled, and precursor level q-value was set to 1%.

#### Differential Expression Testing and Functional Annotation

Approximately five replicate samples were generated for each siRNA and BV2 cell line for a total of 310 samples that were analyzed by mass spectrometry. An average of 6,576 proteins were identified per sample (Figure S2B-C), and some samples had fewer proteins identified with one sample having only 565 proteins identified (Figure S2B-C). The samples with fewer protein IDs tend to also have higher median log2 protein intensity (Figure S2B), which may reflect protein degradation. To limit the potential for these samples to influence downstream analyses, we excluded 11 samples with fewer than 6,100 proteins identified per sample, which resulted in 3 to 6 replicates per siRNA and BV2 cell line (Figure S2A). A large proportion of proteins (4,850) were identified in over 300 samples (Figure S2D). Principal components analysis on the 310 samples (Figure S2E-G) revealed BV2 cell line (i.e., psen2_kd vs scramble) was the biggest contributor to intra-sample variance, and explained 41% of that variance (Figure S2F). Specific target knockdowns contributed less to sample variance, but can be appreciated (Figure S2G). Protein differential abundance was assessed using one-way ANOVA followed by post hoc correction using Tukey’s HSD as previously described [15] (Supplementary Table S4). Gene set enrichment analysis (GSEA) was run with the gseGO function from the clusterProfiler R package (v4.12.6; [16]) against the org.Mm.eg.db annotation database (v3.19.1; [17]). The log2 fold change for each protein was used as a ranking statistic for GSEA and significantly enriched Gene Ontology (GO) terms were mapped onto the TREAT-AD biodomains (syn25428992.v10; [5]) (Supplementary Table S5).

### 2.4 Postmortem Brain Proteomics Data

An integrated meta-analysis of postmortem brain proteomics samples from the AMP-AD consortium was previously performed [5], encompassing 1,188 brain proteomics samples. Gene set enrichment analysis (GSEA) was run with the gseGO function from the clusterProfiler R package (v4.12.6; [16]) against the org.Hs.eg.db annotation database (v3.19.1; [18]). The meta-analysis treatment effect for each protein was used as a ranking statistic for GSEA and significantly enriched Gene Ontology (GO) terms were mapped onto the TREAT-AD biodomains (syn25428992.v10; [5]).

### 2.5 GSEA Term Correlation

Significantly enriched GO terms from the siRNA knockdown proteomics experiments in BV2 cells were compared with the significantly enriched GO terms from AD postmortem brain proteomics. The normalized enrichment score (NES) for GO terms that were either non-significant or not identified in one analysis were set to 0. GO terms were then grouped into biodomains [5], and Kendall’s rank correlation was computed for terms within each biological domain using the NES values from each analysis. The correlation p-values were corrected for multiple testing using Benjamini-Hochberg correction [13].

### 2.6 Biological Domain Protein Correlation

Proteins were grouped into biodomains based on annotation to a term within each domain. For each biological domain, the Pearson correlation was computed between the log fold change values for proteins following target knockdown in BV2 cells and the meta-analysis treatment effect for each orthologous protein from the human dataset. The correlation p-values were corrected for multiple testing using Benjamini-Hochberg correction [13].

### 2.7 Integrating Phenotypic and Proteomic Signatures

The GSEA results for each GO term were compared with the results from the cellular phenotypic assays. For each siRNA and BV2 cell line, we used Spearman’s rho to correlate GO term normalized enrichment scores (NES) with the calculated effect sizes from the linear mixed models for each assay. We did not include the LPS treated NFkB assay results, given that LPS induction was not performed prior to proteomic analysis. The correlation test p-value was corrected for multiple hypothesis testing with Benjamini-Hochberg correction [13]. Significantly correlated terms and phenotype effects are listed (Supplemental Table S6).

### 2.8 Microglia Subtype Datasets

To aid in the characterization of the Psen2 knockdown BV2 cell line, we collected protein lists defining different microglial states and subtypes. Proteomic co-expression modules defined from mouse and human microglial sets were accessed from the publication [19]. Transcripts associated with distinct microglial subtypes were identified from published single cell analyses. Subtype specific gene lists were generated based on supplemental materials from mouse microglial datasets ([20]; filtered FDR < 0.05, homeostatic N = 549, DAM N = 627) and ([21]; filter kME > 0.975, homeostatic N = 396, pro-inflammatory DAM N = 302, anti-inflammatory DAM N = 399). Subtype specific gene lists were also generated from human microglial subtype analyses ([22]; homeostatic N = 178 genes, DAM N = 146 genes) and the gene lists were converted to mouse ortholog identifiers using the gorth function from the gprofiler2 R package (v0.2.3; [23]). GSEA was run against these custom gene lists using the fgseaMultilevel function from the fgsea R package (v1.30.0; [24]). The log fold change values from proteomic assessment of control siRNA treated psen2 knockdown versus control siRNA treated scramble BV2 cells was used as a ranking statistic.

## 3. RESULTS

### 3.1 Psen2 knock-down microglia are pro-inflammatory

The nature of the exemplar hypotheses (i.e. Immune Response and Mitochondrial Metabolism) and subordinate candidate targets selected for exploration suggests that microglial cells would be a relevant cell type to test target function. Microglia are the resident innate immune cells of the brain and influence neuroinflammatory processes that are commonly observed across neurodegenerative conditions, including AD. Moreover, genetic association studies have implicated genes expressed by microglia in AD risk [25,26]. Specific microglial cell states are more strongly implicated in AD pathogenesis [20–22]. To screen for potential AD therapeutic targets it is important to consider cell states that are disease-relevant, and so we chose to use mouse BV2 microglia cell lines in which the Psen2 gene is stably knocked down by shRNA (i.e. psen2_kd) along with a control line (i.e. scramble). Mutations in both PSEN1 and PSEN2 are causal for autosomal dominant forms of AD [27] and in bulk transcriptomic meta-analyses of AMP-AD data the PSEN2 transcript is significantly decreased in AD brains relative to control brains (Figure S3A). It has been shown that Psen2 functions are distinct from Psen1 functions in microglia and that either γ-secretase inhibition or specifically Psen2 knockdown in these BV2 cells results in an exaggerated cytokine response [9].

When we compared the function of the Psen2 knockdown (psen2_kd) BV2 cell line with the scramble BV2 cell line we noted an array of phenotype differences that indicate the Psen2 knockdown cells are a relevant model for AD microglial states. The Psen2 knockdown condition affects the expression of nominated target genes (Figure S1C), and in particular Ifih1 which is 37-fold up-regulated in the Psen2 knockdown cells. When we compare the proteomes between Psen2 knockdown and scramble cell lines, there is increased expression of proteins from a number of GO terms within the Immune Response, Apoptosis, Lipid Metabolism and Oxidative Stress biodomains, among others (Figure 2A). The up-regulation of Immune Response, Lipid Metabolism, and Oxidative Stress domain terms are significantly positively correlated with terms that are impacted in AD (Figure 2B). Specifically, of the 197 Immune Response domain terms that are up in AD postmortem brain, 60 are also up in the Psen2 knockdown BV2 cells when compared with the scramble BV2 cells (Figure 2C). Along with up-regulated expression of Immune Response biodomain proteins, we also noted a higher baseline of positivity from the phagocytosis assays in the Psen2 knockdown cell line compared with the scramble cells (e.g., see Figure 3A). We tested microglial proteomic co-expression modules [19] and found that the proteins up-regulated in the Psen2 knockdown cells are significantly enriched for a module related to “Macrophage cytokine production”, while proteins that are down-regulated in Psen2 knockdown cells are significantly enriched for an “RNA processing” module (Figure S3B). Finally, when we tested for proteins associated with different transcriptomically defined microglial subtypes [20–22], we found that an enrichment for disease associated and homeostatic microglial subtypes indicating that proteins associated with these relevant microglial cell states are expressed higher in Psen2 knockdown BV2 cells than the scramble cells. In summary, the Psen2 knockdown BV2 cell line is relatively more pro-inflammatory and may represent a cell state that is more disease-relevant than the control BV2 cells.

**Figure 2.**
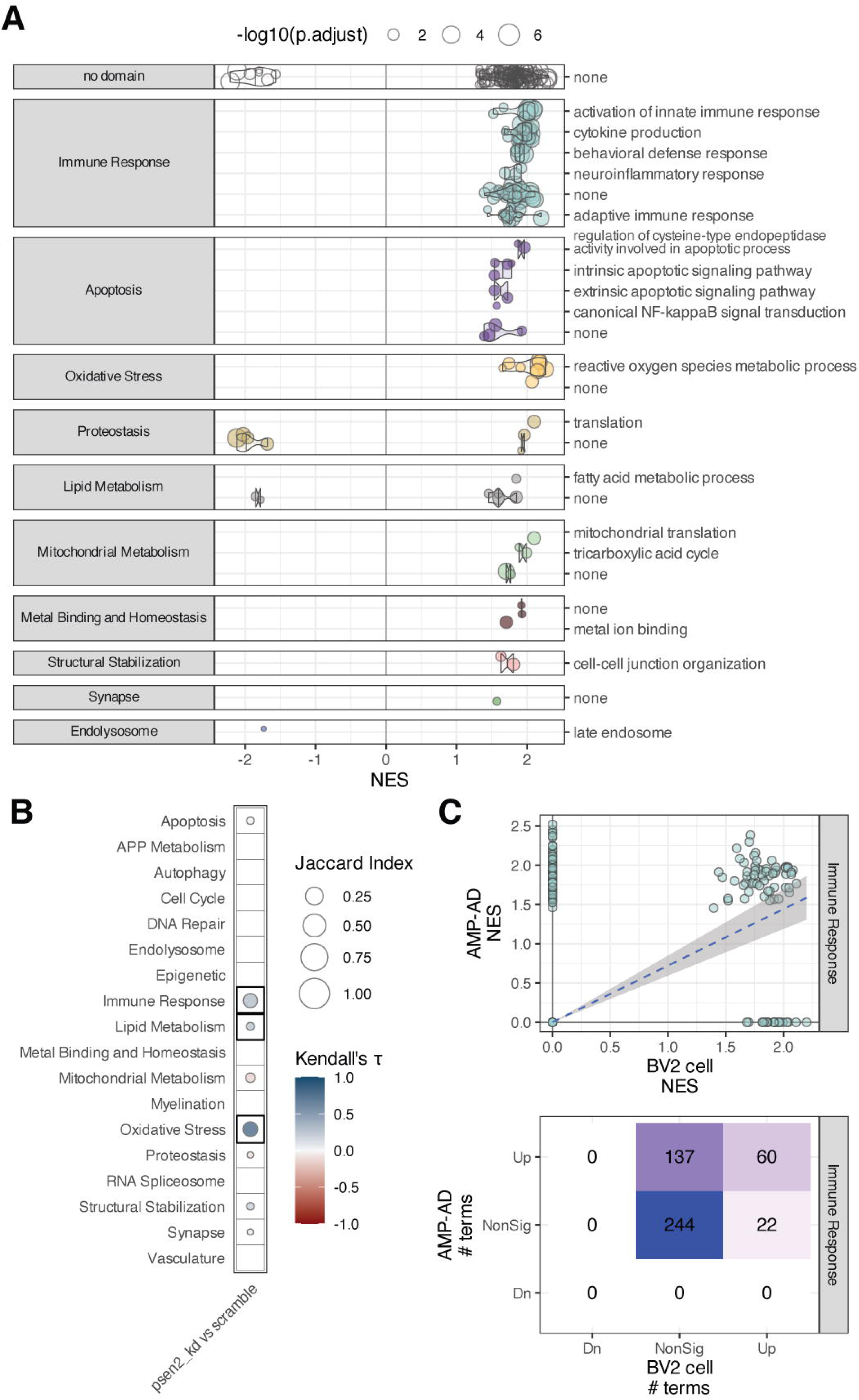
Proteomic assessment of Psen2 knockdown BV2 microglia compared with wild-type (i.e., scramble) BV2 microglia. (A) GSEA results showing significantly differentially expressed GO terms (points) from each biological domain (facets) and how the proteins from those terms are differentially expressed in Psen2 knockdown BV2 cells (x-axis). The size of each point corresponds to the -log_10_ adjusted p-value for the term enrichment. (B) Correlation plot across AD biodomains showing domains where GO term enrichments are correlated with enrichments from late-onset AD brain proteomics from AMP-AD. The size of each point reflects the size of the overlap between analyses and the fill color indicates the correlation direction. Correlations that are significant following multiple hypothesis testing correction are outlined with a box. (C) Example of Immune Response biological domain terms enriched in each case showing the distribution of term NES values from each analysis (top) and the numbers of terms in each class (bottom).

**Figure 3.**
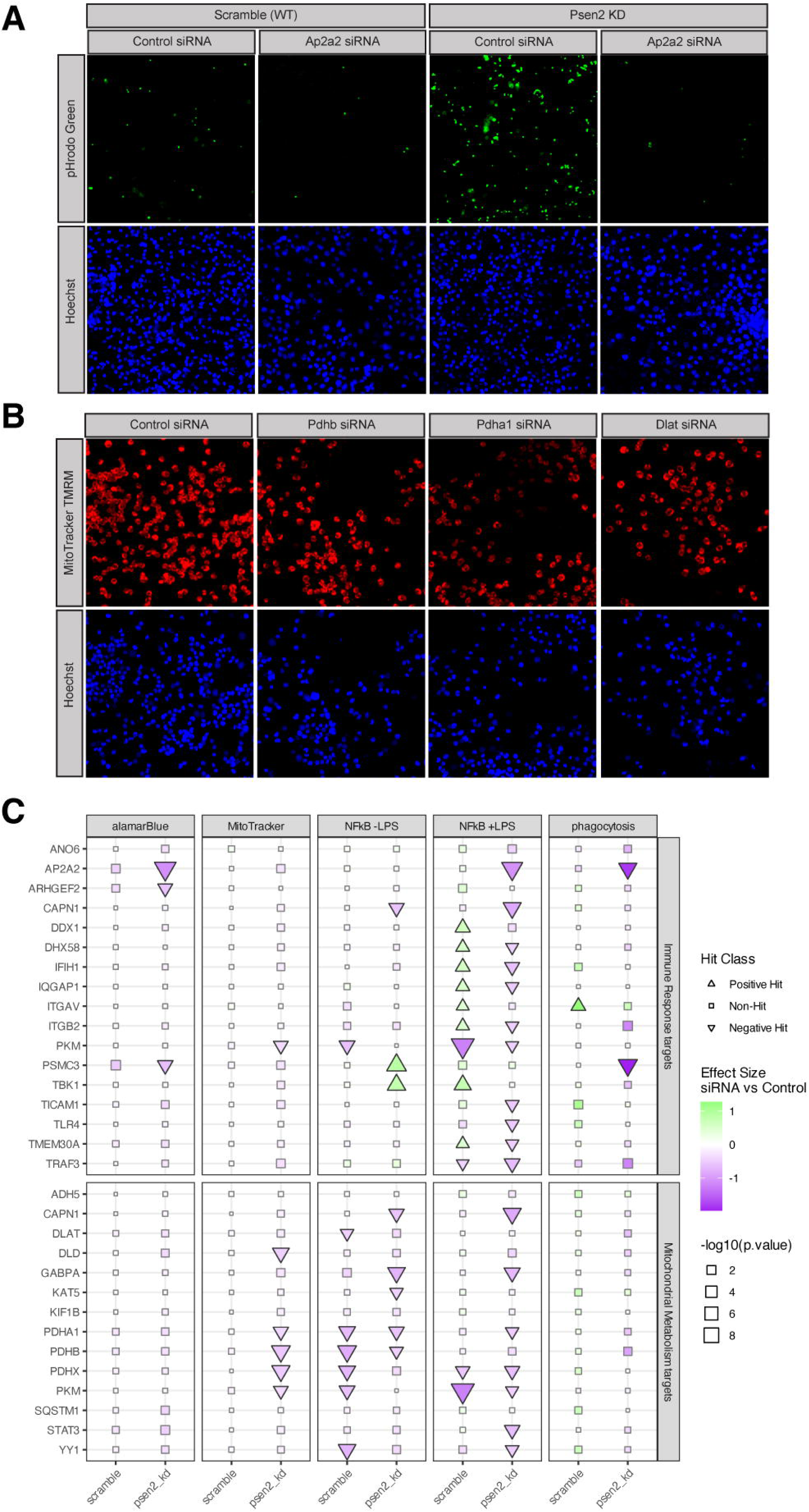
BV2 cell phenotypes following target knockdown. (A) Example images from the pHrodo Green phagocytosis assay (top) and nuclear counter-stain (bottom) from control siRNA and Ap2a2 siRNA treated cultures. The results from both wild-type and Psen2 knockdown cell lines are shown. (B) Example images from the MitoTracker TMRM assay (top) and nuclear counter-stain (bottom) in Psen2 knockdown cells from control siRNA and three different target knockdowns (Pdhb, Pdha1, and Dlat). (C) Summary of all assay results for each target. Targets that modify the indicated phenotype with an adjusted p-value ≤ 0.05 are plotted with a triangle, the size of each point is scaled to the significance of the observed effect and the fill for each point shows the computed effect size of target knockdown in the indicated assay.

### 3.2 Target perturbations impact relevant cellular phenotypes

We validated the knockdown of targets by siRNA in BV2 cells using real-time PCR assays prior to further testing of cellular phenotypes and proteomic responses. We confirmed over 50% knockdown for most targets, with many as high as 70-80% knockdown (Figure S1C). Two targets had knockdown levels below 50%; TICAM1 knockdown averaged a 46% reduction in expression while DHX58 knockdown averaged a 19% reduction (Figure S1C).

Cellular phenotype screening was performed following 72 hours of siRNA knockdown of targets in each BV2 cell line. Example images are shown for the phagocytosis assay (Figure 3A) and MitoTracker assay (Figure 3B). Because of the baseline differences between the Psen2 knockdown and scramble cell lines, especially for the immune assays, the cell lines were analyzed separately and results are shown separately for each background. A summary of all results from cell phenotype assays is shown (Figure 3C) and more detailed results are contained in the supplemental figures (alamarBlue, Figure S4; MitoTracker, Figure S5; NFkB Figure S6; phagocytosis Figure S7). Overall, 25 of the 29 targets (86% hit rate) were identified as a significant hit in at least one assay across our panel. The assays showed differential sensitivities to perturbation, with the NFkB reporter assay following LPS induction having the highest hit rate (19 hits, 65%), followed by NFkB reporter without LPS induction (11 hits, 38%), MitoTracker (5 hits, 17%), and phagocytosis and alamarBlue (3 hits each, 10%). Importantly, the Psen2 knockdown cells were more sensitive to target knockdown. For the mitochondrial phenotypes (i.e. alamarBlue and Mitotracker assays), all hits were exclusive to the Psen2 knockdown cells. For the immune phenotypes there were more hits in the Psen2 knockdown than scramble cells; for the phagocytosis assay there were 2 hits in Psen2 knockdown and 1 in scramble, and for the NFkB reporter following LPS induction there were 16 hits for Psen2 knockdown cells and 11 hits for scramble. For the NFkB reporter assay in the absence of LPS induction there were an equivalent number of hits in each cell line (7 hits), though 5 hits were unique for one cell line or the other and only 2 hits were common between cell lines. There were also 5 hits from the NFkB reporter assay following LPS induction that had different directionality in the different cell lines. The hits for the MitoTracker assay were specific to targets nominated from the mitochondrial hypothesis area, while hits for the phagocytosis and alamarBlue assays were specific to targets nominated from the immune hypothesis area, and NFkB hits were from both hypothesis areas. The most impactful target perturbations include Ap2a2, Psmc3, Itgav, Pkm, Pdha1, Pdhb, and Pdhx which were each hits in multiple different assays. These results highlight the impact of perturbing the nominated targets on relevant cellular phenotypes and emphasize the importance of the disease-relevant Psen2 knockdown cell line for sensitization to those perturbations.

### 3.3 Identification of targets that reverse disease associated proteomic signatures

As with the cellular phenotypes, the proteomic analyses of target knockdowns revealed a range in the degree of impact. Most targets had very few significantly differentially expressed proteins identified, while a few had hundreds to over a thousand significant protein differences (Figure 4A).

**Figure 4.**
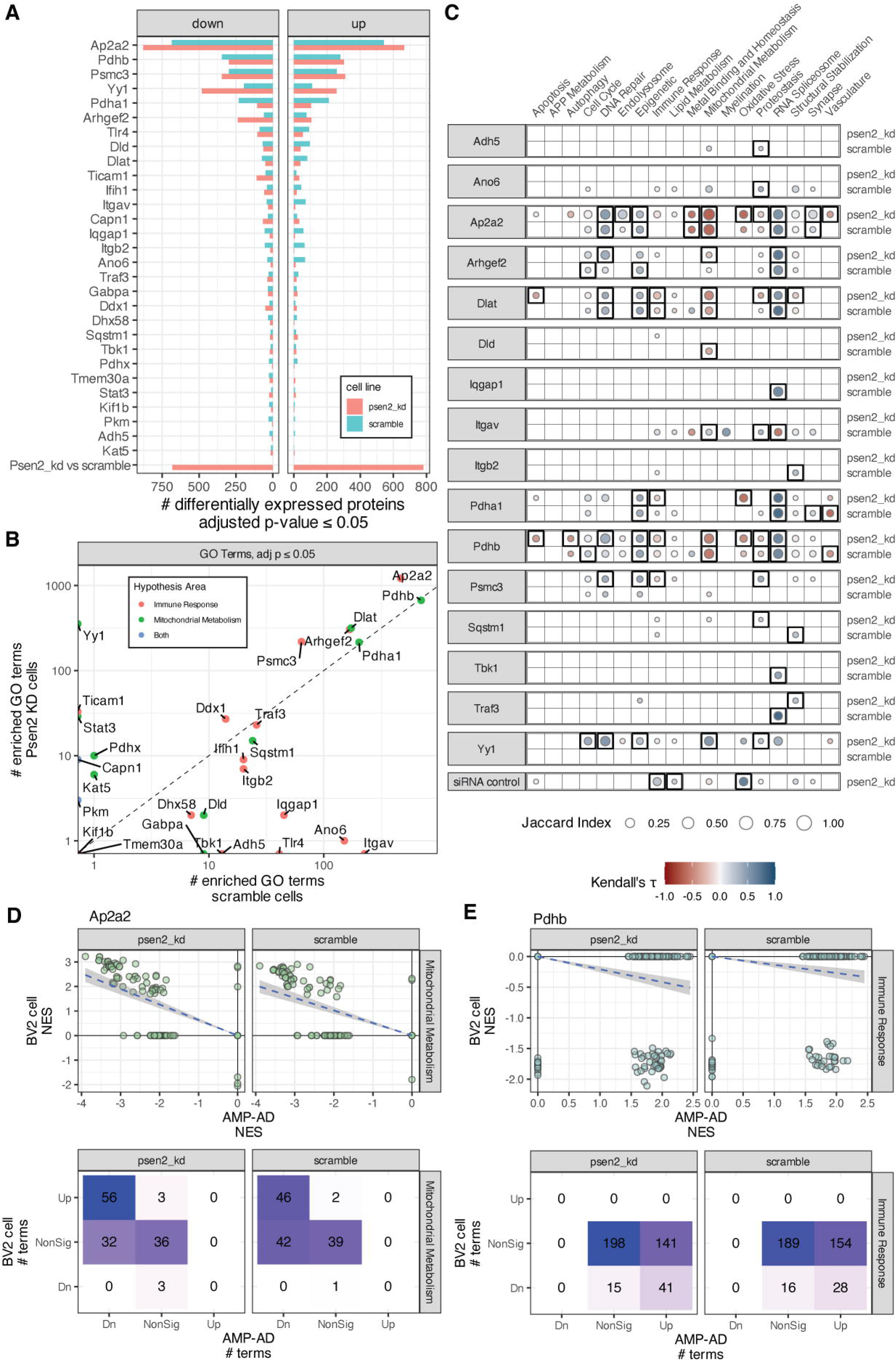
Proteomic responses to target knockdown. (A) The number of significantly (adjusted p-value ≤ 0.05) differentially expressed proteins in each target knockdown and each BV2 cell line. (B) The number of significantly enriched GO terms from GSEA analyses for each target in each BV2 cell line. (C) Correlation plots across AD biodomains, targets, and cell lines showing domains where GO term enrichments are correlated with enrichments from late-onset AD brain proteomics from AMP-AD. The size of each point reflects the size of the overlap between analyses and the fill color indicates the correlation direction. Correlations that are significant following correction for multiple hypothesis testing are outlined with a box. (D) Example of Mitochondrial Metabolism domain term enrichments from AMP-AD and Ap2a2 knockdown in each cell line showing the distribution of term NES values from each analysis (top) and the numbers of terms in each class (bottom). (E) Example of Immune Response domain term enrichments from AMP-AD and Pdhb knockdown in each cell line showing the distribution of term NES values from each analysis (top) and the numbers of terms in each class (bottom).

We examined the proteomic effect on the target of each knockdown and found that for all but five proteins (Ifih1, Traf3, Tlr4, Ticam1, and Kat5) there was a negative log fold change for the targeted protein (Figure S2H). The perturbation with the largest impact was Ap2a2, with 913 significantly affected proteins in scramble cells and 1,292 significantly affected proteins in Psen2 knockdown cells. Yy1, Ap2a2, and Arhgef2 each had several hundred more differentially expressed proteins in Psen2 knockdown cells than scramble cells, while Pdha1 had more differentially expressed proteins in scramble cells. GSEA tests were run to identify GO terms that were significantly altered within each perturbation experiment (Figure 4B). The targets with the largest number of differentially expressed proteins (i.e. Ap2a2, Pdhb, Yy1, etc) also had the largest number of significantly enriched GO terms. Most targets had a similar number of enriched terms from each cell line, but as with the differential expression analysis Yy1 knockdown had a larger impact on Psen2 knockdown cells with 361 significantly enriched terms from Psen2 knockdown cells and none from scramble cells. Conversely, Itgav and Ano6 had over 140 significantly enriched terms in scramble cells and either 0 or 1 from Psen2 knockdown cells.

Biological domain enriched terms were correlated between target knockdowns in BV2 cells and postmortem brain proteomics from AD cases versus controls (Figure 4C). 16 of the 29 targets tested had biological domain term enrichments that were significantly correlated with AD. Of these, Pdhb, Dlat, Ap2a2, Pdha1, Yy1, and Psmc3 had significant correlations over four or more biodomains. Several knockdowns induce term enrichments that are anti-correlated with AD. Knockdown of Ap2a2, Arhgef2, Dlat, Dld, and Pdhb each resulted in increased expression of proteins annotated to Mitochondrial Metabolism GO terms, reversing the broad down-regulation of these functions that are associated with AD in postmortem brain proteomics. For example, knockdown of Ap2a2 resulted in the up-regulation of proteins from 46 and 56 GO terms from the Mitochondrial Metabolism domain of the 88 that are down in AD (Figure 4D). Protein level correlation by biological domain supports the reversal of Mitochondrial Metabolism disease associated signatures by knockdown of Ap2a2, Arhgef2, Dlat, Dld, and Pdhb (Figure S8). Similarly, knockdown of Dlat, Pdha1, Pdhb, and Psmc3 each resulted in decreased expression of proteins annotated to Immune Response GO terms, reversing the up-regulation of these processes that is seen in AD brain proteomics. For example, knockdown of Pdhb resulted in the down-regulation of 28 and 41 GO terms from the Immune Response domain of the 182 that are up in AD (Figure 4E). All of the effects of Yy1, which are restricted to Psen2 knockdown cells, result in changes that are positively correlated with AD. There are also positive correlations with disease for many of these knockdowns in the DNA Repair, Epigenetic and RNA Spliceosome domains. Overall, there is strong support for perturbation of Ap2a2, Dlat, and Pdhb inducing a reversal of disease-associated proteomic signatures in BV2 cells.

### 3.4 Integrated cell phenotypes and proteomics identifies top-tier targets

Alignment between the cell phenotype and proteomics highlighted relevant GO term enrichment patterns that were correlated with phenotypic effect sizes (Figure 5A). Of the GO terms that are significantly correlated with the MitoTracker effect sizes, the most directly relevant is “mitochondrial inner membrane”, the NES values for which are negatively correlated with assay effect size (Spearman’s ρ = −0.4, adjusted p = 0.016). This suggests a compensatory response where knockdowns that lead to decreases in MitoTracker signal tend to also drive an up-regulation in the expression of mitochondrial inner membrane proteins. Likewise the alamarBlue assay effect sizes were negatively correlated with NES values for the “regulation of mitotic cell cycle phase transition” GO term (Spearman’s ρ = −0.679, adjusted p = 8.4×10^-6^). This suggested that perturbations leading to a decrease in metabolically active cells are also increasing the expression of proteins that regulate cell cycle checkpoints and arresting cell division. Conversely, the immune assays tend to have positive correlations with relevant terms. For example, the effect sizes from the phagocytosis assay are positively correlated with the NES values for the “phagocytosis” GO term (Spearman’s ρ = 0.468, adjusted p = 0.0097), which reveals a direct mechanistic link where the increase in cellular phagocytosis is associated with the increased expression of the proteins involved in the process. The effect sizes for the NFkB assay are also positively correlated with the NES values for the GO term “proteasome accessory complex” (Spearman’s ρ = 0.519, adjusted p = 0.0299), which is consistent with the role of the proteasome in the activation of NFkB signaling through destruction of IkB [28]. By linking phenotypic outcomes with their corresponding molecular correlates, this approach enables the identification of targets that exert coordinated effects on both cellular phenotypes and the biological processes captured by GO terms.

**Figure 5.**
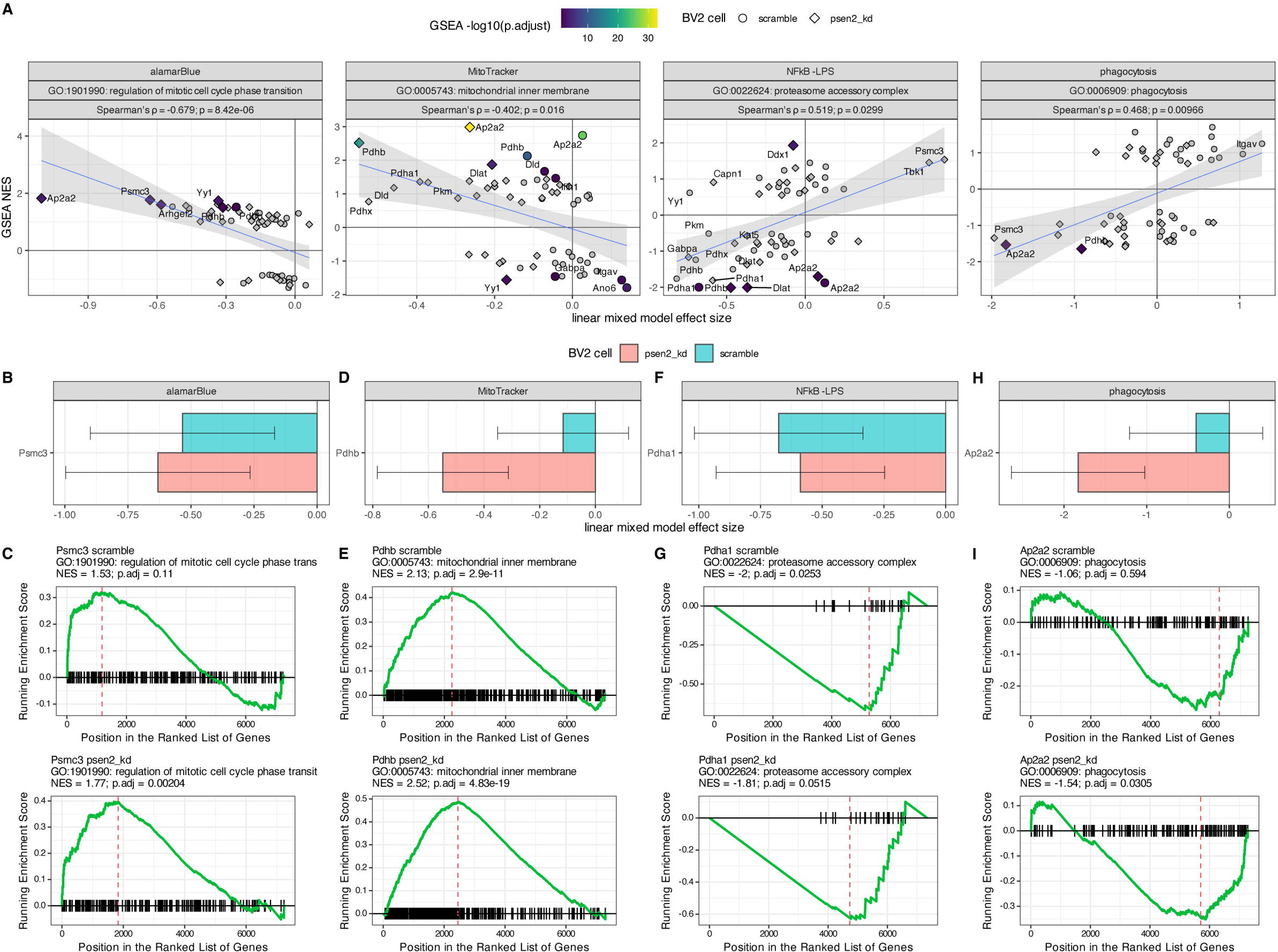
Integration of cell phenotype effects and GO term enrichments for target knockdowns. (A) Correlation of cellular assay effect size (x-axis) with GO term NES value (y-axis) for the assay and GO term indicated in the header of each plot. The header also shows the correlation statistics between the assay effect size and term enrichment values. The shape of each point denotes the corresponding cell line and knockdowns that resulted in a significant term enrichment have a fill value corresponding to the significance of the term enrichment. Detailed examples of specific assay results are shown for Psmc3 (B), Pdhb (D), Pdha1 (F) and Ap2a2 (H). Detailed examples of GSEA term enrichment statistics for specific GO terms are shown for Psmc3 (C), Pdhb (E), Pdha1 (G), and Ap2a2 (I).

Specific examples of these associations are shown (Figure 5B-I). Psmc3 inhibition leads to a decrease in alamarBlue signal in both scramble and Psen2 knockdown cell lines, with a slightly stronger effect in the Psen2 knockdown line (Figure 5B). This is mirrored in the GSEA enrichment patterns for “regulation of mitotic cell cycle phase transition”, which has a larger NES value and a smaller p-value for the inhibition of Psmc3 in the Psen2 knockdown cell line (Figure 5C). The knockdown of Pdhb has a much stronger effect on the MitoTracker assay readout in Psen2 knockdown cells (Figure 5D). In this case the proteomic response for the “mitochondrial inner membrane” is strong and significant for both cell lines, though the NES is larger and the enrichment has a smaller p-value in the Psen2 knockdown cell line (Figure 5E). The inhibition of Pdha1 has a stronger effect on NFkB reporter output in the scramble cell line (Figure 5F), and the relative strength is reflected in the enrichment statistics for the “proteasome accessory complex” GO term which has a more negative NES value and a smaller p-value in the scramble cell line (Figure 5G). The knockdown of Ap2a2 has a larger impact on the phagocytic activity of Psen2 knockdown cell line than the scramble cell line (Figure 5H), which is reflected in the GSEA statistics for the “phagocytosis” term following Ap2a2 knockdown, which has a negative NES value but is not significant in the scramble cell line and has a more negative NES value and is significant in the Psen2 knockdown cell line.

Taken together, the identification of specific targets that significantly alter both cellular phenotypes and the related molecular signatures provides compelling evidence that these perturbations directly impact the underlying biology.

## 4. DISCUSSION

Effective prioritization of candidate targets is challenging but essential for advancing novel AD therapeutics. We have developed an experimental framework to assess prioritized targets in a cell-specific context. The screening platform established here involves perturbation of nominated targets in an appropriate cell line, and subsequent assays of cellular functions relevant to the hypotheses under investigations along with measurement of molecular changes. The molecular readouts also enable direct comparison with disease omic measures to highlight which target perturbations drive molecular changes that are correlated or anti-correlated with disease associated molecular changes. Through integration of the results from assays of cellular phenotypes with the specific and related proteomic changes, we identify the most robust targets that have direct impacts on the hypothesized biological functions and represent the strongest candidates for further development efforts.

We screened 29 candidates linked to immune or mitochondrial hypotheses in BV2 microglia, and measured phenotypes and proteomic responses. We used this hypothesis validation workflow to filter the list of 29 candidate targets to five top-tier targets that both impact relevant phenotypes and induce proteomic changes that reverse AD-associated signatures: AP2A2, PDHB, PDHA1, DLAT, and PSMC3. Three of these targets (PDHA1, PDHB, and DLAT) encode protein components of the E1 and E2 subunits of the pyruvate dehydrogenase (PDH) complex, PSMC3 is a component of the 19S regulatory subunit of the proteasome, and AP2A2 is the alpha subunit of the adaptor protein complex AP-2.

Decreased expression and function of genes and proteins involved in mitochondrial bioenergetics, including PDH subunits, is an early and consistent hallmark of AD associated neurodegeneration (Figure S1A, [29]). Animal model studies supports the role of PDH complex members in AD pathophysiology including that PDH kinase inhibition prevented neuron loss and improved memory performance [30], that conditional knockout of Pdha1 impairs memory function [31], and Dlat knockdown reduced neuronal damage and cognitive deficits [32]. There are likely cell-type specific implications of modulation of PDH given that phosphorylation levels of PDH are correlated with neuronal activity [33], while shifts between oxidative phosphorylation and glycolysis are associated with changes in microglial states [34]. The results from our experiments support inhibition of PDH subunits impacting both mitochondrial membrane polarity and NFkB signaling (Figure 3C, Figure 5A), as well as reversing disease associated proteomic signatures in both the Mitochondrial Metabolism and Immune Response biodomains (Figure 4C). Both PDHA1 and PDHB are targets that have been nominated by several AMP-AD teams for therapeutic development [35] and here we find further evidence supporting their therapeutic potential.

Like the PDH complex, there is extensive evidence that proteasome function is impaired early in AD pathogenesis [36,37], and impaired proteasome functions are associated with memory deficits in mice [38]. While various neurodegeneration-associated oligomeric proteins, including Aβ, directly inhibit proteasome function [39,40], increased proteasome activity has been observed in reactive glia around Aβ plaques [41]. While the proteasome is involved in activating NFkB signalling [28], it also influences microglial cytokine signaling and memory performance in mouse models [42]. The specific implication of the PSMC3 subunit is supported by genetic fine mapping that identified PSMC3 among others at the CELF1/SPI1 locus [43] and methylation QTL studies that implicate differential methylation at the locus in AD risk [44]. Inhibition of Psmc3 induced a reversal of disease related Immune Response biological domain signatures and directly impacted immune and cell viability phenotypes, especially in the disease relevant Psen2 knockdown BV2 microglia line (Figure 3-5).

Given the proteasomal involvement in activation of NFkB signaling, it is notable that inhibition of Psmc3 resulted in increased NFkB reporter activity. However Psmc3 inhibition also resulted in generally increased expression of proteasome accessory subunit proteins, particularly in Psen2 knockdown BV2 cells (Figure 5A). Overall these results support the therapeutic potential for modulation of the proteasome, including PSMC3, in AD.

The AP2 complex participates in the assembly of clathrin-coated vesicles (CCV) at the plasma membrane and induces clathrin mediated endocytosis along with CCV accessory proteins and AD risk genes PICALM and BIN1 [45]. AP2 interacts with PICALM and LC3 to mediate the endocytosis and autophagy of the APP C-terminal fragment [46], and regulates the endosomal trafficking and lysosomal delivery of BACE1, influencing amyloid generation [47]. AP2 subunits have been found in detergent insoluble fractions in AD brains [48], further supporting the association of the AP2 complex with neuropathological accumulation. Rare variation in a variable number tandem repeat (VNTR) region near AP2A2 is associated with neocortical phospho-tau burden [49]. Importantly, the AP2 alpha subunits differentially associate with intraneuronal paired helical filaments (AP2A1) and IBA1^+^ microglia (AP2A2) in human brain tissue [48], suggesting an approach to drive interventions toward either microglial or neuronal AP2 complexes. This is an important consideration given that neuronal specific knock-outs of AP2 complex members lead to increased Aβ production [47], whereas our results endorse inhibition of Ap2a2 in microglia as therapeutically beneficial. Ap2a2 knockdown in the Psen2 knockdown BV2 cell line reduced both microglial phagocytosis and NFkB signaling (Figure 3C), as well as decreased cell viability, perhaps in conjunction with alterations of cell cycle checkpoints (Figure 5A). Inhibition of Ap2a2 in BV2 microglia also reversed disease-associated proteomic signatures in the Mitochondrial Metabolism and Metal Binding and Homeostasis domains, and reversed Oxidative Stress, Proteostasis, and Vasculature biological domain signatures specifically in the Psen2 knockdown cells (Figure 4C). AP2 subunits including AP2A1, AP2B1, and AP2M1 have each been nominated by AMP-AD investigators for therapeutic development [35] and here we provide evidence supporting the therapeutic potential of an additional AP2 subunit, AP2A2.

An important throughline in our data is the importance of including a sensitized cell condition. Using Psen2 knockdown BV2 cells revealed heightened baseline phagocytosis and AD-relevant proteomic signatures (Figure 2, Figure 3, Figure S2). Inclusion of the Psen2 knockdown line gets us closer to understanding the effects of target perturbation in a disease-relevant context. Indeed our results highlight many phenotypic and proteomic responses that are significant only in the sensitized cell line. While the inclusion of Psen2 knockdown cells here is revelatory, future efforts will explore additional disease relevant stressors such as exposure to oligomeric Aβ or oligomeric tau that may be more directly relevant to more common, late-onset forms of AD.

We found limited support for hypothesis specificity of target perturbations, potentially indicated coordinated disease relevance of metabolic and immune dysfunction. For example, only targets nominated from the Mitochondrial Metabolism hypothesis area impacted MitoTracker assay phenotypes and only targets nominated from the Immune Response hypothesis area had significant effects in the phagocytosis assay (Figure 3C). However, only Immune Response targets affected the metabolically sensitive alamarBlue assay phenotype, and both target sets affected NFkB reporter phenotypes (Figure 3C). Moreover, the proteomic responses of the targets showed no hypothesis specificity as the Mitochondrial hypothesis targets Pdhb, Pdha1 and Dlat all reversed Immune Response domain proteomic responses, whereas the Immune hypothesis targets Ap2a2 and Arhgef2 both reversed Mitochondrial Metabolism domain proteomic responses (Figure 4C). In the context of immune response and mitochondrial functions this is perhaps unsurprising, given what is known about the interplay of bioenergetics and immune functions [50–52]. We would not have appreciated this lack of target specificity were it not for our unbiased, multi-dimensional screening strategy. We intend to refine the target and hypothesis nomination process based on the lessons learned here and, for example, would focus future hypotheses on points of functional convergence between biodomains.

Several other limitations of this work restrict the nature of the conclusions we are able to draw. Because our screening system relies on immortalized murine BV2 cells, which diverge from human microglia *in vivo*, the results of this work should be viewed as primarily a filter for the nominated targets. We’re able to rapidly prioritize candidates for further study using these results, and definitive target validation will require testing in human iPSC derived microglia or *in vivo* models. Another limitation is the reliance on siRNA-mediated target knockdown, which introduces variability in the levels of target suppression (e.g. Figure S1C) and may include off-target effects that confound interpretation. Follow-up with approaches such as CRISPRa/CRISPRi systems which enable the activation or inhibition, respectively, of target genes would provide orthogonal data to support our conclusions. Indeed the results from some targets imply that increasing expression of the target may be preferable — e.g. knockdown of Yy1 produced disease correlated effects across domains suggesting that perhaps activation would lead to anti-correlated, therapeutic effects.

### 4.1 Conclusion

We have developed a robust platform for screening TREAT-AD nominated targets for functional engagement of biological hypotheses. Employing this strategy filtered the list of 29 candidate targets to five top-tier targets that are promising leads for further investigation and resource development by the Emory-Sage-SGC-Jax TREAT-AD center. The results from this work, as well as all enabling reagents developed through work by the center will be made openly available as a resource to the AD research community. Ultimately, the approach developed here demonstrates potential to accelerate AD therapeutic target discovery.

## Abbreviations

AD: Alzheimer’s Disease
TREAT-AD: Target Enablement to Accelerate Therapy Development in AD
AMP-AD: Accelerating Medical Partnership for AD
GSEA: Gene Set Enrichment Analysis
GO: Gene Ontology

## ACKNOWLEDGEMENTS

The authors would like to thank Dr. Suman Jayadev for providing the BV2 cell lines used in this work. Data used in this study were obtained from the Accelerating Medicines Partnership Program for Alzheimer’s Disease (AMP AD) Consortium members below: Mayo RNAseq Study: Study data were provided by the following sources: The Mayo Clinic Alzheimer’s Disease Genetic Studies, led by Dr. Nilufer Ertekin Taner and Dr. Steven G. Younkin, Mayo Clinic, Jacksonville, FL, using samples from the Mayo Clinic Study of Aging, the Mayo Clinic Alzheimer’s Disease Research Center, and the Mayo Clinic Brain Bank. Data collection was supported through funding by NIA grants P50 AG016574, R01 AG032990, U01 AG046139, R01 AG018023, U01 AG006576, U01 AG006786, R01 AG025711, R01 AG017216, R01 AG003949, NINDS grant R01 NS080820, CurePSP Foundation, and support from Mayo Foundation. Study data include samples collected through the Sun Health Research Institute Brain and Body Donation Program of Sun City, Arizona. The Brain and Body Donation Program is supported by the National Institute of Neurological Disorders and Stroke (U24 NS072026 National Brain and Tissue Resource for Parkinson’s Disease and Related Disorders), the NIA (P30 AG19610 Arizona Alzheimer’s Disease Core Center), the Arizona Department of Health Services (contract 211002, Arizona Alzheimer’s Research Center), the Arizona Biomedical Research Commission (contracts 4001, 0011, 05 901, and 1001 to the Arizona Parkinson’s Disease Consortium), and the Michael J. Fox Foundation for Parkinson’s Research. Religious Orders Study/Memory and Aging Project (ROSMAP): We are grateful to the participants in the Religious Order Study and the Memory and Aging Project. This work was supported by the US National Institutes of Health (U01 AG046152, R01 AG043617, R01 AG042210, R01 AG036042, R01 AG036836, R01 AG032990, R01 AG18023, RC2 AG036547, P50 AG016574, U01 ES017155, KL2 RR024151, K25 AG041906 01, R01 AG30146, P30 AG10161, R01 AG17917, R01 AG15819, K08 AG034290, P30 AG10161, and R01 AG11101). Mount Sinai Brain Bank (MSBB): This work was supported by grants R01AG046170, RF1AG054014, RF1AG057440, and R01AG057907 from the NIH/NIA. R01AG046170 is a component of the AMP AD Target Discovery and Preclinical Validation Project. Brain tissue collection and characterization was supported by NIH HHSN271201300031C.

## CONSENT STATEMENT

No human participants were recruited for this work. The details of the Institutional Review Board (IRB)/oversight body that provided approval or exemption for the research described are given as follows: Western Institutional Review Board—Copernicus Group (WCG) IRB of Sage Bionetworks gave ethical approval for this work.

