## Supplemental Figures for "Integrated phenotypic and proteomic screening identifies top-tier Alzheimer’s disease therapeutic targets"

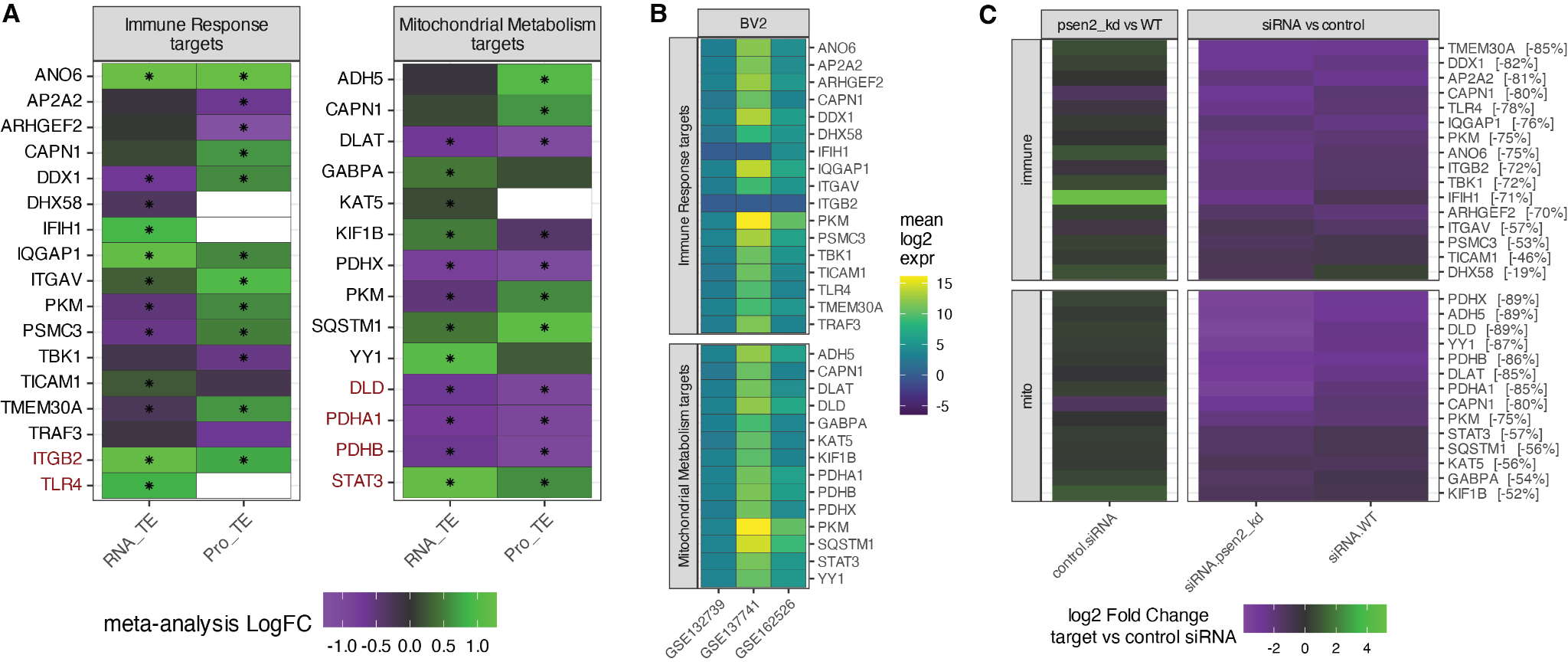

**Figure S1.** Nominated target details and knockdown validation. (A) Heatmap of differential transcript (RNA_TE) and protein (Pro_TE) abundance in AD versus control brains post mortem from our meta-analysis of AMP-AD data. Significant differences are indicated with an asterisk. Targets that have been previously nominated by AMP-AD investigators are indicated in red. (B) Heatmap of target expression levels in BV2 cell data from GEO datasets. (C) Heatmap showing real-time PCR results of target expression comparing Psen2 knockdown versus wild-type cells (left) and following shRNA knockdown of each target (right). For each target knockdown, the average change in target expression between target shRNA and control shRNA are shown as a percentage difference in brackets along the y-axis.

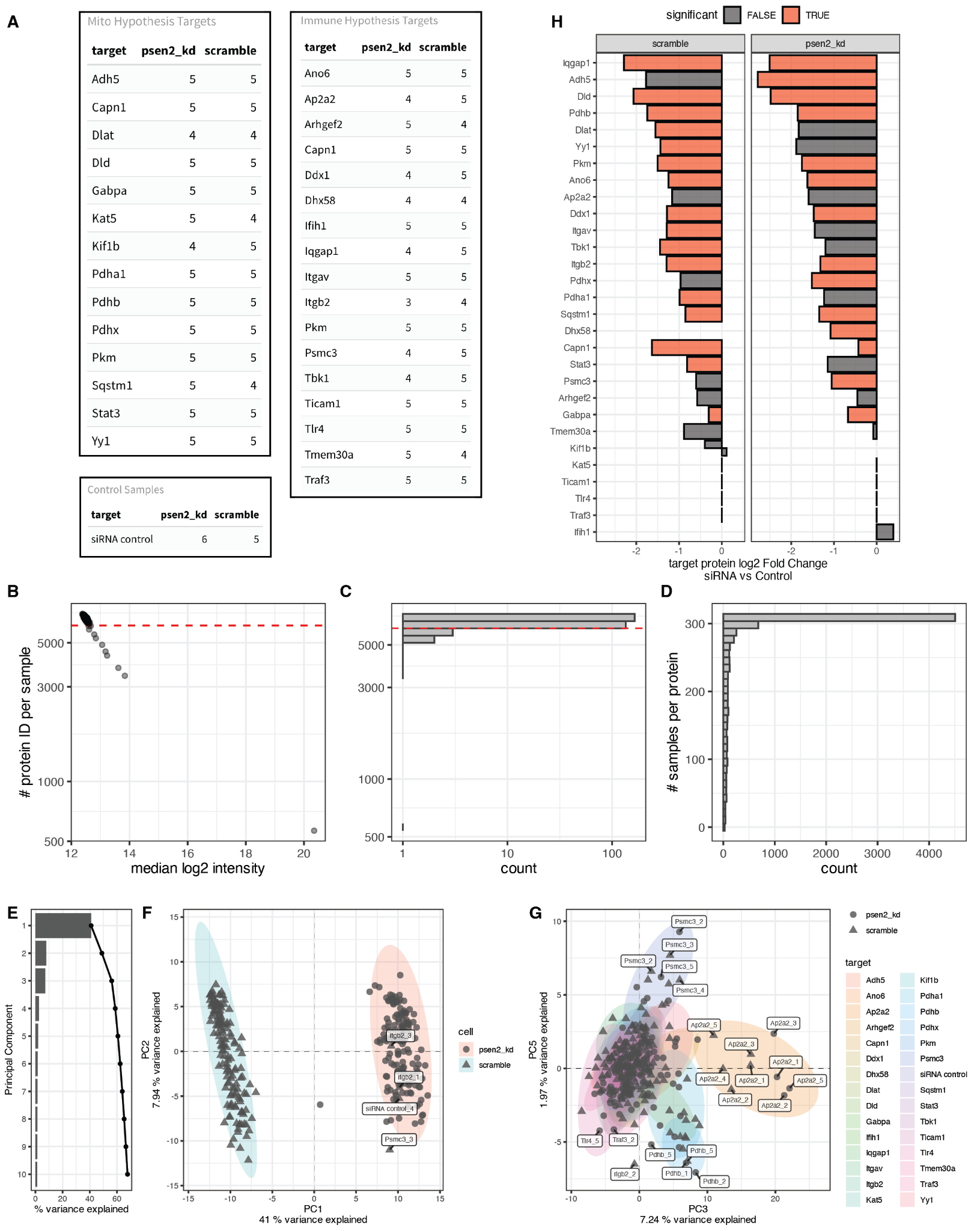

**Figure S2.** Proteomic assay quality control. (A) The number of samples for each condition following sample quality control. (B) The relationship between the number of proteins ID in each sample (y-axis) and the median log2 intensity of proteins with the sample (x-axis). The dashed line shows the location of 6100 proteins per sample which was used to remove samples from the analysis. (C) Histogram showing the distribution of the number of proteins per sample along the y-axis in panel B. The dashed line shows the location of 6100 proteins per sample which was used to remove samples from the analysis. (D) Histogram showing the number of samples in which each protein was identified. (E) Principal component eigenvalues for the first 10 components. (F) Scatterplot of principal components 1 and 2. The shape of each point denotes the cell line of each sample either Psen2 knockdown cells (circle) or scramble cells (triangle) and ellipses represent a 95% CI around the centroid for each cell line group, Psen2 knockdown (red) and scramble (blue). Individual scramble samples that are grouped with Psen2 knockdown are labelled. (G) Scatterplot of principal components 3 and 5. Point shapes correspond to the cell line, either Psen2 knockdown (circle) or scramble (triangle), while ellipses represent a 95% CI around the samples from a group defined by a common siRNA target. (H) Proteomic assessment of target knockdown showing the log2 fold change for each protein in the corresponding knockdown experiment along the x-axis. The fill of each bar represents whether the identified difference was found to be significant in a given experiment (adjusted p-value ≤ 0.05, red) or not.

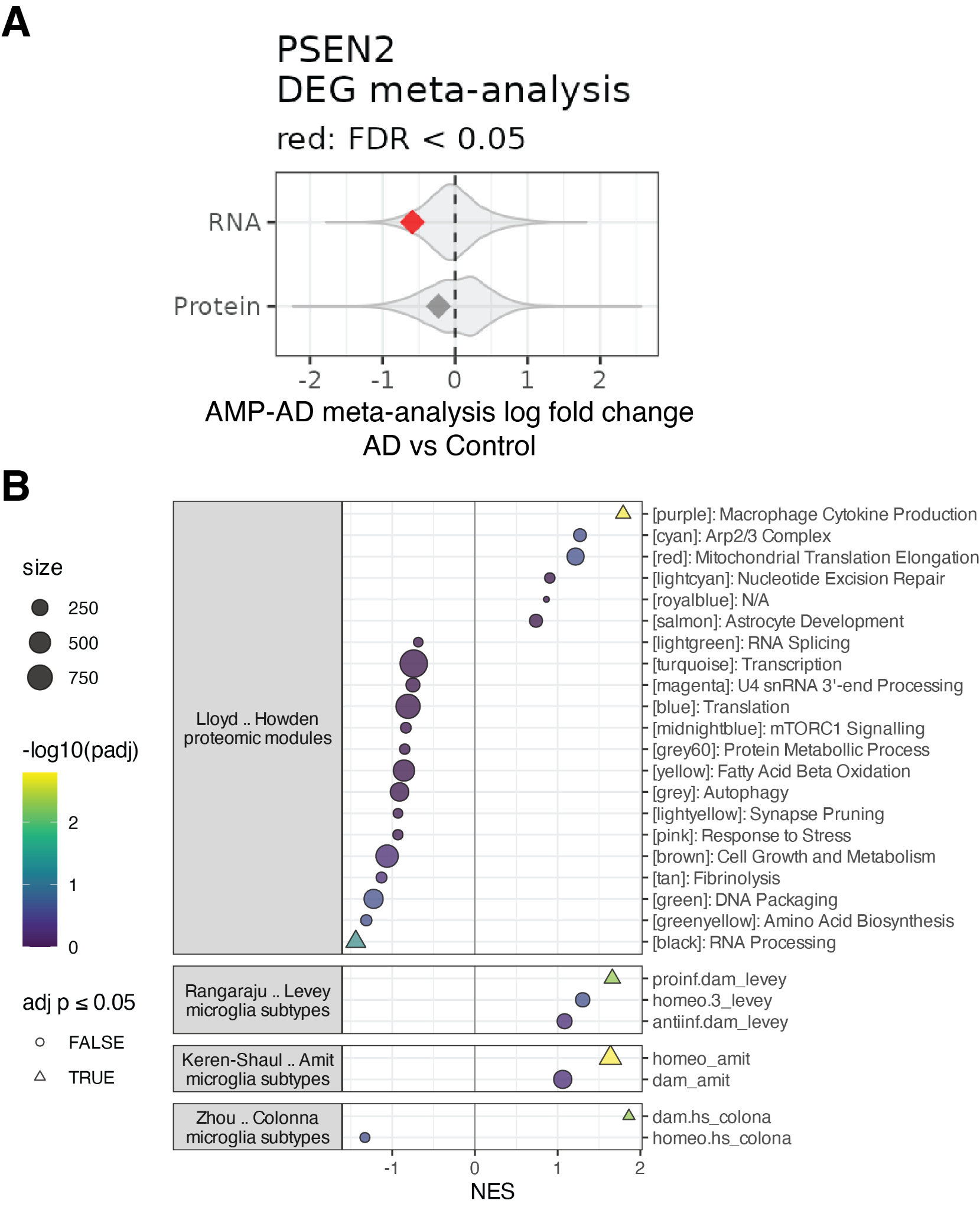

**Figure S3.** Relevance of Psen2 knockdown microglia to AD. (A) Summary of AMP-AD meta-analyses. Violin plots show the distribution of transcriptomic (RNA) and proteomic (Protein) treatment effects and the diamond points show the specific values for the Psen2 transcript and protein. Psen2 RNA is significantly lower in AD brains relative to controls, and Psen2 protein is lower, though it is not significant. (B) GSEA values for microglial gene sets as defined in methods. The NES values for all sets are shown with the size of each point corresponding to the number of genes in each set and the fill color corresponding to the significance of the enrichment. Sets that are significantly enriched (adjusted p-value ≤ 0.05) are plotted as a triangle.

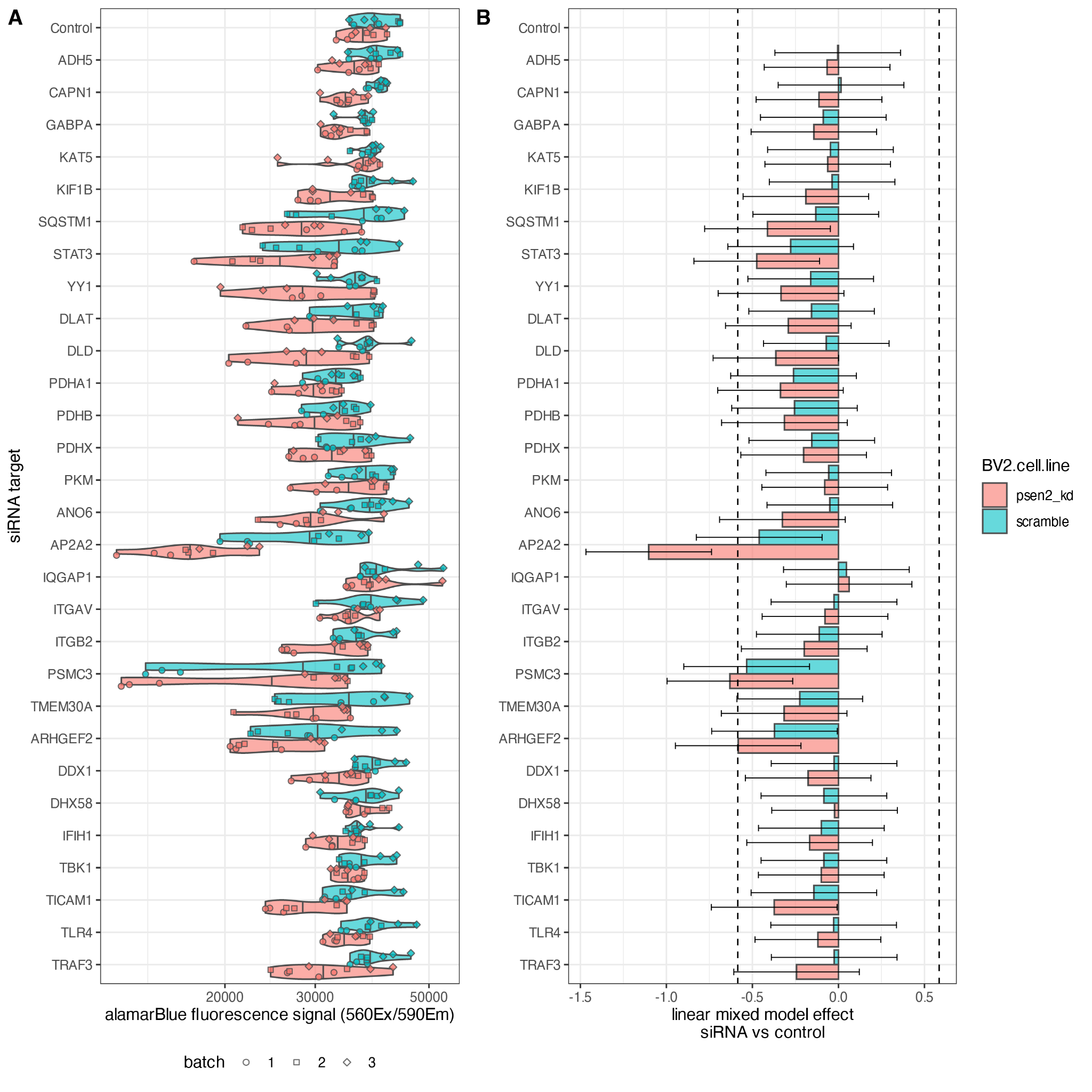

**Figure S4.** Alamar Blue assay results. (A) Raw alamarBlue fluorescence signal (Ex^560^/Em^590^) for each sample measured. Samples point colors correspond to the BV2 cell line and shapes correspond to the experimental batch for each measurement. (B) Summary of linear mixed model results showing the effect size for each target in each BV2 cell line. Error bars show the model confidence interval. Dashed lines show levels that correspond to 50% increase or decrease.

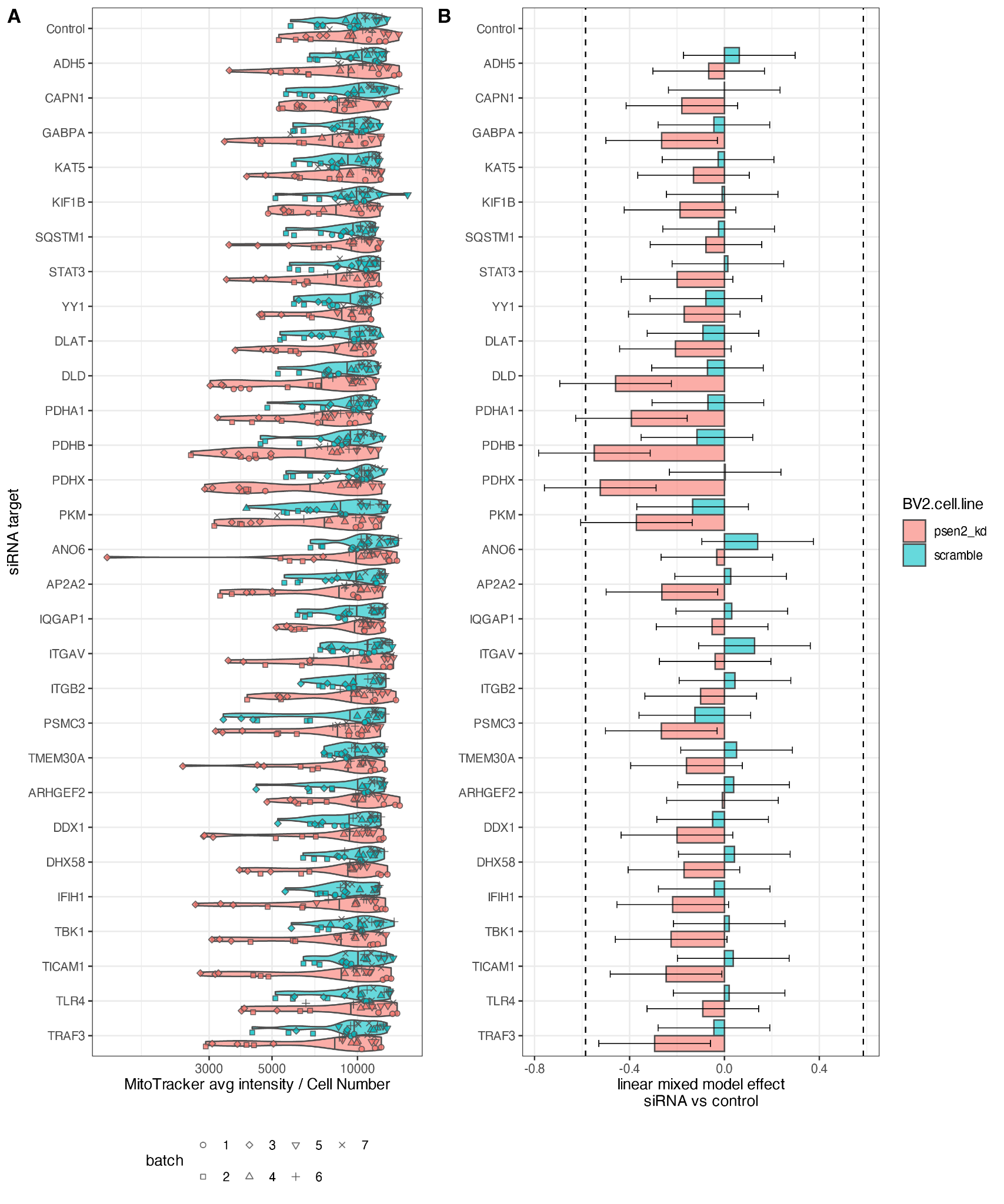

**Figure S5.** MitoTracker TMRM assay results. (A) Raw MitoTracker average fluorescence intensity normalized to cell number for each sample measured. Samples point colors correspond to the BV2 cell line and shapes correspond to the experimental batch for each measurement. (B) Summary of linear mixed model results showing the effect size for each target in each BV2 cell line. Error bars show the model confidence interval. Dashed lines show levels that correspond to 50% increase or decrease.

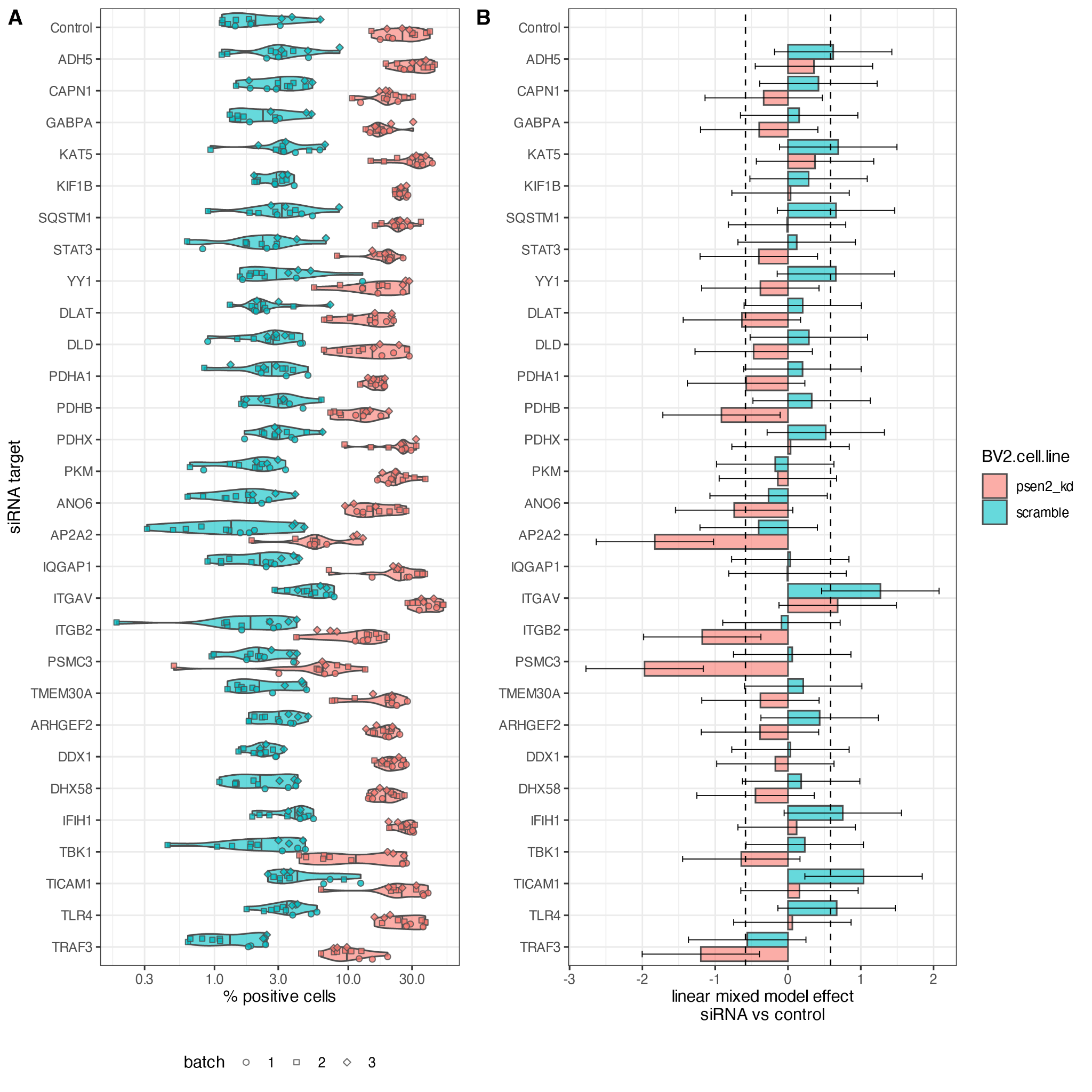

**Figure S6.** pHrodo Green assay results. (A) Raw percent of cells positive for pHrodo green for each sample measured. Samples point colors correspond to the BV2 cell line and shapes correspond to the experimental batch for each measurement. (B) Summary of linear mixed model results showing the effect size for each target in each BV2 cell line. Error bars show the model confidence interval. Dashed lines show levels that correspond to 50% increase or decrease.

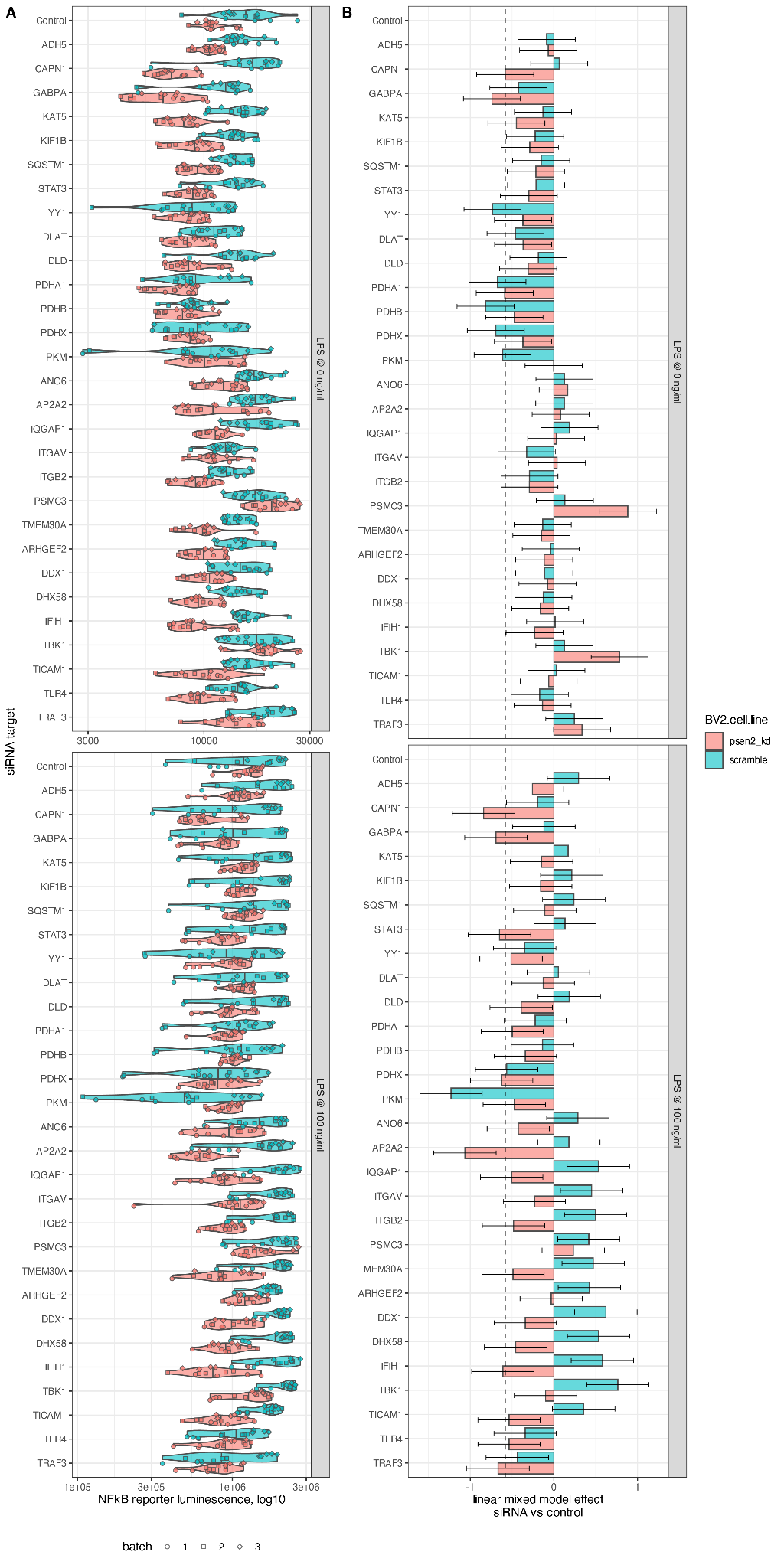

**Figure S7.** NFkB reporter assay results. (A) Raw NFkB reporter luminescence intensity on a log10 scale for each sample measured. Samples point colors correspond to the BV2 cell line and shapes correspond to the experimental batch for each measurement. The values for samples treated with 100 ng/ml LPS are shown (bottom) along with those that received no LPS induction (top). (B) Summary of linear mixed model results showing the effect size for each target in each BV2 cell line. Error bars show the model confidence interval. Dashed lines show levels that correspond to 50% increase or decrease. The values for samples treated with 100 ng/ml LPS are shown (bottom) along with those that received no LPS induction (top).

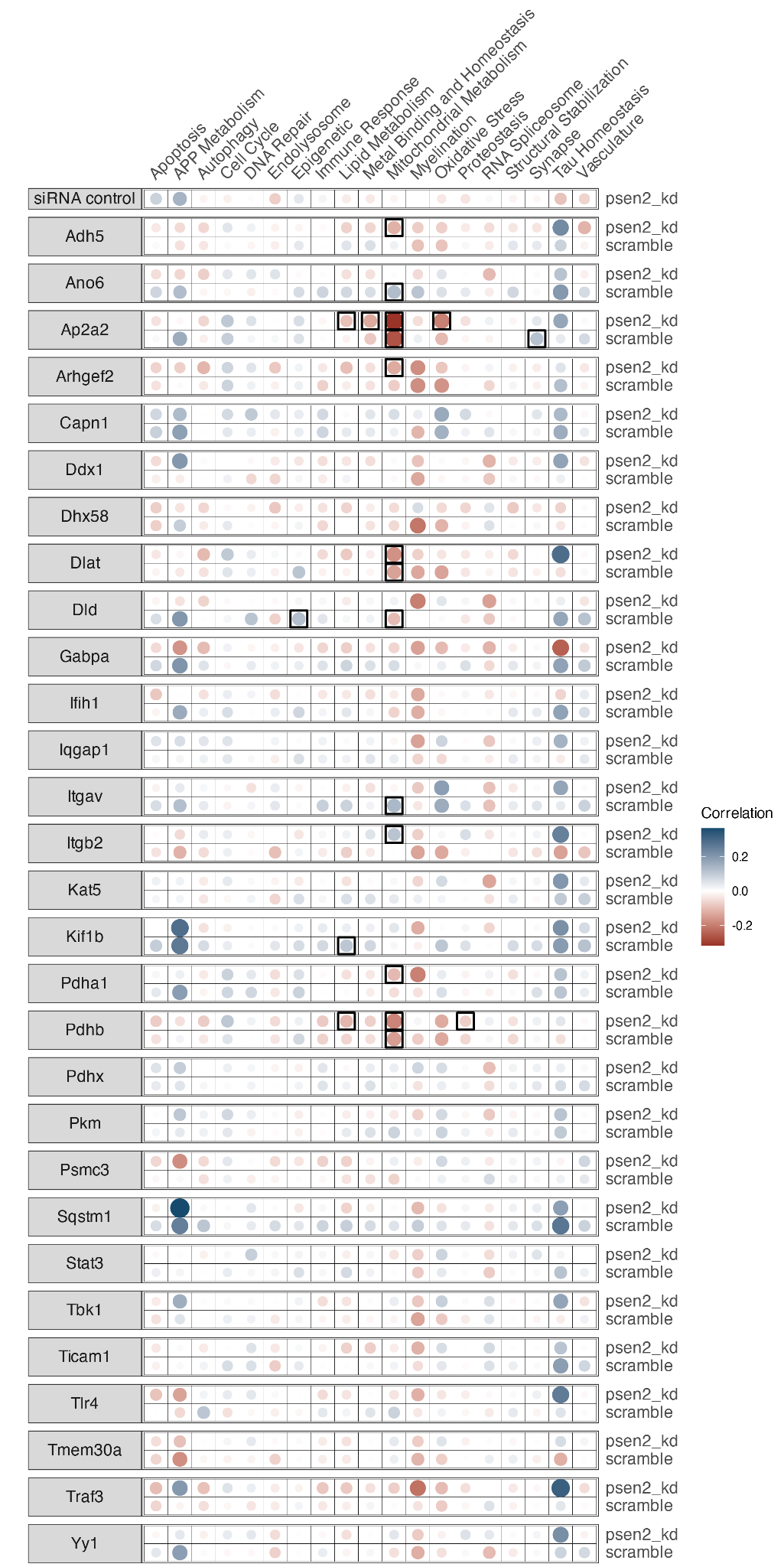

**Figure S8.** Correlation of protein log2 fold change values between shRNA knockdown in BV2 cells and orthologous proteins from AMP-AD proteomics. Protein groups were defined by AD biodomain annotation. Pearson correlation was used to relate the change in proteins within each group between the BV2 cell experiments (i.e., target shRNA vs control shRNA) and AMP-AD proteomics (i.e., AD vs control). The fill color of each point represents the correlation coefficient for each comparison and the size of each point corresponds to the absolute value of the computed correlation coefficient. Any correlation that was found to be significant after correction for multiple hypothesis testing is outlined with a box.
